# Disentangling Schwann Cell and Neuronal TRPA1 Function in Mouse Models of Familial Episodic Pain Syndrome

**DOI:** 10.64898/2026.02.14.705765

**Authors:** Matilde Marini, Martina Chieca, Elisabetta Coppi, Marco Montini, Lorenzo Bonacchi, Lorenzo Landini, Irene Scuffi, Kelvin Y Kwan, Alice Papini, Gaetano De Siena, Elisa Bellantoni, Lucrezia Timotei, Valentina Albanese, Gabriele Ferroni, Evelynn Dalila do Nascimento Melo, Marie-Christine Birling, Romain Lorentz, Romina Nassini, Francesco De Logu

## Abstract

Familial Episodic Pain Syndrome (FEPS) is a rare inherited disorder characterized by episodes of severe upper-body pain triggered by different stimuli including cold, stress, or fasting. A gain-of-function point mutation (N855S) in the Transient Receptor Potential Ankyrin 1 (TRPA1) ion channel has been identified in affected individuals, altering its biophysical properties, and leading to sustained nociceptive signaling. While TRPA1 is predominantly studied in sensory neurons, recent findings highlight its key modulatory role for Schwann cells in chronic pain. Here, we investigated the cell-specific contributions of mutant TRPA1 (TRPA1*) in FEPS by developing mouse models with TRPA1* selectively expressed in either Schwann cells or sensory neurons, using CRISPR-based and Cre-loxP strategies. Patch-clamp analyses confirmed that TRPA1* exhibits enhanced current responses to agonists compared to wild-type. Through behavioral assays we revealed that TRPA1* expressed in sensory neurons mediates acute nociception, while TRPA1* in Schwann cells drives mechanical allodynia in response to subthreshold doses of TRPA1 agonists and to physiological pain triggers commonly observed in FEPS patients, including fasting, cold exposure, and restraint stress. Pain responses were associated with the increase in reactive oxygen species (ROS) and accumulation of 4-hydroxynonenal (4-HNE) in TRPA1* sciatic nerves and these effects were reduced by a treatment with an antioxidant. We reveal distinct roles of neuronal and non-neuronal TRPA1 in pain and provide novel *in vivo* models to investigate the mechanisms of chronic pain in FEPS and related channelopathies. Overall, this study offers new insights into the development of targeted therapies for Schwann cell-TRPA1 to relieve pain in affected individuals.

## Introduction

Inherited mutations in ion channels can disrupt normal ion flux across the plasma membrane, leading to a range of conditions, including pathological pain^1,2^. Non-functional mutations in voltage-gated sodium channels (Nav) have been linked to a lack of pain sensation in congenital indifference to pain syndrome^3–5^ and to autosomal dominant familial hemiplegic migraine with aura^6–8^. Conversely, missense mutations in Nav channels have been associated with severe pain episodes, such as those observed in primary erythromelalgia syndrome^9^. Similarly, mutations in voltage-gated calcium channels (Cav) have been linked to conditions such as trigeminal neuralgia^10^ and to susceptibility to malignant hyperthermia^11^.

The Transient Receptor Potential (TRP) family of channels includes multimodal ion channels responding to a wide range of physical and chemical stimuli, including changes in temperature, mechanical pressure, and various chemical irritants^12–16^. Mutations in several TRP channels have been correlated with distinct channelopathies, including TRPC6 with focal segmental glomerulosclerosis^17^, TRPV4 with autosomal dominant metatropic dysplasia and Charcot-Marie-Tooth disease type 2C^18,19^, TRPM1 with autosomal recessive congenital stationary night blindness^20^, TRPML1 with mucolipidosis type IV^21^, and TRPP2 with autosomal dominant polycystic kidney disease^22^. However, the only known inherited human pain-associated TRP channelopathy involves a mutation in the TRPA1 subtype and is called Familial Episodic Pain Syndrome (FEPS)^23^. A gain-of-function point mutation (N855S) in the S4 domain of TRPA1 was first identified in a South American family. The altered biophysical properties caused by TRPA1 channel mutation resulted in episodes of severe pain in the upper body triggered by conditions of food restriction, cold exposure, and stress. Moreover, it enhances the sensitization of the nociceptive system as demonstrated by exaggerated and prolonged hyperalgesia and allodynia following application of the TRPA1-selective agonist mustard oil^24^.

Conventionally, pain has been attributed to the activation/sensitization of sensory neurons, which detect noxious stimuli and transmit a pain signal to the central nervous system. However, recent discoveries highlight the complex interaction between neuronal and non-neuronal cells in pain development and maintenance^25–31^. Schwann cells, known for their role in protecting and supporting peripheral nerves, have recently been recognized for their role in mediating mechanical and pressure sensation in the skin^32–34^ and in development and maintenance of mechanical allodynia in different mouse models of pain^35–43^. Here, by comparing the wild-type (TRPA1) and mutated (TRPA1*) mouse ion channels we confirmed that the FEPS gain-of-function point mutation altered the channel biophysical properties, with the TRPA1* channel exhibiting larger current than the outward-rectifying WT channel. In addition, using three distinct genetic approaches we developed a robust FEPS mouse model with cell-selective expression of TRPA1* in sensory neurons (Adv-TRPA1* mice) or Schwann cells (Plp-TRPA1*mice), in which we identified distinct roles for TRPA1 in mediating acute nociception and mechanical allodynia, respectively. To dissect the role of neuronal and non-neuronal cells in FEPS, we also used three behavioural procedures in mice (food deprivation, stress induction, and cold temperature exposure) that reflect the primary triggers of body pain in FEPS patients. We reported that *Plp*-TRPA1* but not *Adv-*TRPA1* mice were sensitive to the development of mechanical allodynia following fasting, stress, and cold-temperature exposure. The feed-forward mechanism activated by TRPA1* in Schwann cells by a subthreshold dose of reactive oxygen species (ROS) facilitated and sustained mechanical allodynia in *Plp-*TRPA1*.

Our study provides evidence that neuronal and non-neuronal TRPA1 mediates distinct mechanisms of pain perception in mouse models of FEPS, with neuronal TRPA1 primarily mediating acute nociception. At the same time, Schwann cell TRPA1 is essential for sustaining mechanical allodynia. Overall, the presented FEPS mouse model provides a valuable tool for investigating the distinct roles of neuronal and Schwann cells in pain pathways, supporting the development of targeted therapeutic strategies for chronic pain.

## Results

### Conservation of the N855S mutation and biophysical characterization of mouse TRPA1*

FEPS is caused by an A to G transition in exon 22 of the *TRPA1* cDNA (c.A2564G) which results in the substitution of an asparagine by a serine (N855S) in the transmembrane segment S4^24^. To establish a relevant mouse model for in vivo functional studies, we first confirmed that the human mutation site region was conserved in the mouse Trpa1 (Fig. 1a). We next characterized the biophysical properties of the WT (TRPA1) and mutated mouse TRPA1 (TRPA1*) channels. Plasmids encoding either TRPA1 or TRPA1* were expressed in HEK293T cells and analyzed using whole-cell patch-clamp recordings. Consistent with findings in human TRPA1^44^, the overall voltage-dependent currents activated by a depolarizing voltage ramp protocol (−100 to +100 mV) in TRPA1 and TRPA1* cells were identical in absence of TRPA1 agonists (Fig. 1b). However, upon stimulation with selective exogenous or endogenous TRPA1 agonists, allyl isothiocyanate (AITC) or 4-hydroxynonenal (4-HNE) respectively (both 100 µM; 2 min application), TRPA1* channel mediated significantly larger currents compared with the TRPA1 wild type counterpart (Fig. 1c-j). Indeed, averaged TRPA1*-activated currents, obtained by subtraction of the control ramp from that recorded in the presence of AITC (Fig. 1e) or 4-HNE (Fig. 1i), were at least doubled in the outward range up to +100 mV whereas the inward component, negligible in TRPA1-transfected cells upon agonist exposure (Fig. 1e and i), was prominent. Both channels were inhibited by the selective TRPA1 antagonist A967079 (Fig 1 c,d,g,h). These results indicate that the conserved N858S mutation in mouse TRPA1 reproduces the gain-of-function effect described in human TRPA1^44^, consistent with FEPS pathogenesis and provide a basis for generating mouse models of the disease.

**Figure 1.**
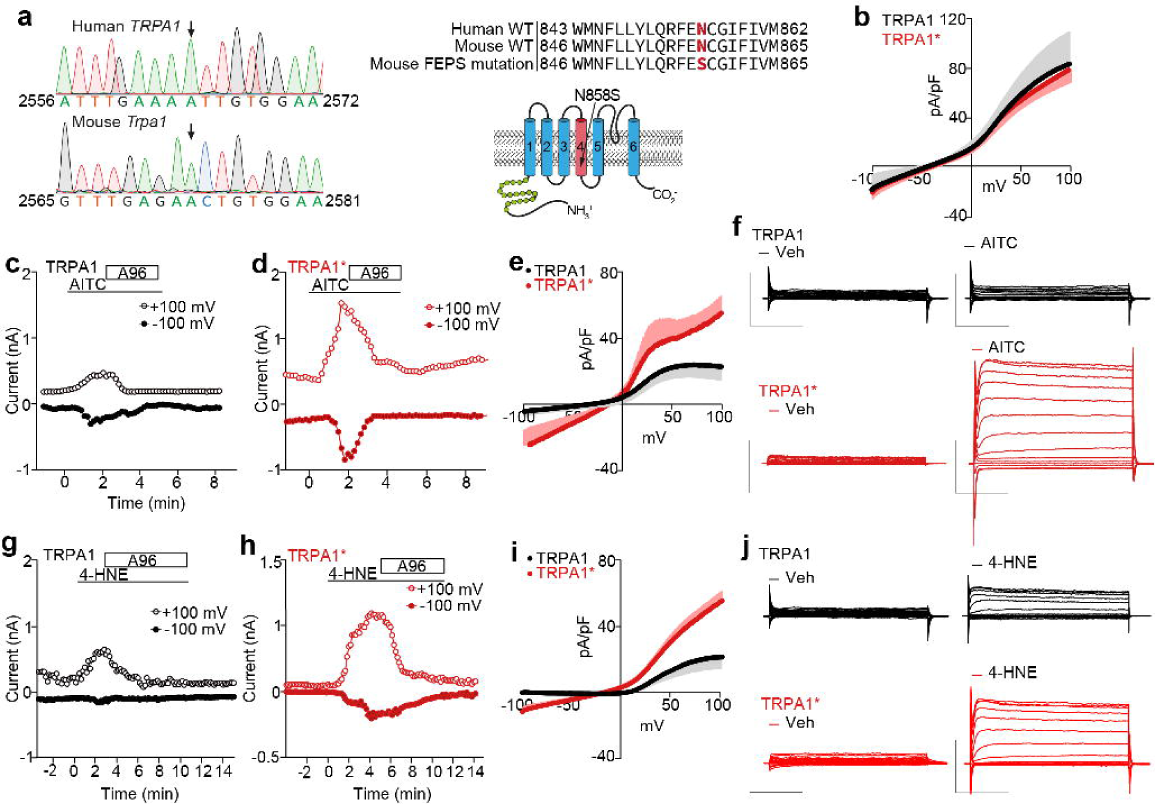
HEK293 cells transfected with the mutant mTRPA1* channel present enhanced currents upon stimulation with exogenous or endogenous agonists. **(a)** Sanger sequence chromatogram showing human *TRPA1* and mouse *Trpa1*. Amino acid sequences are conserved between *wild type* (WT) human and mouse genes. Schematic representation of FEPS mutation (N858S) in transmembrane domain 4 of TRPA1 channel. **(b)** Voltage-dependent currents activated a depolarizing voltage ramp protocol (from −100 to +100 mV; 800 ms) in HEK293 cells transfected with the mouse TRPA1 channel (TRPA1, n=12) or its mutated counterpart (TRPA1*, n=15). **(c,d)** Time course of current amplitude evoked by the voltage ramp at + 100 mV or −100 mV in a TRPA1- (**c**) or TRPA1*- (**d**) HEK293T transfected cell after AITC (100 µM) also in the presence of A967079 (A96) (100 µM). **(e)** AITC-activated currents measured in TRPA1-(n=6) or in TRPA1*-(n=6) HEK293T transfected cells by subtraction of the control ramp from that recorded in AITC. **(f)** Original current traces evoked by a depolarizing voltage-step protocol (from −140 to +140 mV; 20 mV increment; 300 ms steps) recorded before or after the application of AITC in a TRPA1 or TRPA1*-HEK293T transfected cell **(g,h)** Time course of current amplitude evoked by the voltage ramp at + 100 mV or −100 mV in a TRPA1-(**c**) or TRPA1*- (**d**) transfected HEK293 cell after 4-HNE (100 µM) and in the presence of A96 (100 µM). **(i)** 4-HNE-activated currents measured in TRPA1-(n=6) or in TRPA1*-(n=9) HEK293T transfected cells. **(f)**. Original current traces evoked by a depolarizing voltage-step protocol recorded before or after the application of 4-HNE in a TRPA1- or TRPA1*-HEK293T transfected cell. Data are mean ± s.e.m.

### Generation of mouse models carrying the TRPA1* mutation

We recently observed that the Schwann cell (SC) and neuronal TRPA1 channel play a critical and distinct role in several mouse model of pain including neuropathic and inflammatory pain, cancer and migraine pain^36,37,39,41,42,45,46^. Here we used the gain-of-function TRPA1 channel with altered biophysical properties to examine the roles of SC- and neuronal-mediated TRPA1 in a mouse model of FEPS. To investigate the physio/pathological role of the TRPA1* channel in pain, we used an adeno-associated viral (AAV) vector-based strategy to supplement functionally mutated copies of the wild-type *Trpa1* gene in mice. We developed two complementary approaches, both based on a double AAV platform that combines CRISPR-Cas9 genome editing with micro-homology-mediated end-joining (MMEJ)^47^ to achieve targeted expression of *Trpa1* in vivo* in mice. Both platforms involve AAV-mediated, Cre-dependent expression of a loxP-flanked *Staphylococcus aureus* Cas9 (SaCas9) and eGFP (Fig. 2a,b). In the first strategy, termed genome-editing approach (GEdit), SaCas9 and two gRNAs excised the endogenous *Trpa1* sequence from the mouse genome of *Cre*-expressing mice and the exogenous *Trpa1\** sequence from the AAV vector (*Plp*-*Cre*-GEdit mice for Schwann cells and *Adv*-Cre-GEdit mice for sensory neurons) (Fig. 2a). AAVrh10 (intravenous administration) or AAVPHP.S (intrathecal administration) were used in *Plp*-Cre or *Adv*-Cre mice respectively^41,48^. The *Trpa1\** sequence was then integrated in the genome using microhomology arms (MHA) with single-base-pair substitutions introduced at the gRNA recognition sites to prevent repeated cleavage (Fig. 2a). The second strategy, termed minigene approach (MiniG), was developed as a complementary method. Here, we used Cre-*Trpa1^fl/fl^* mice generated by crossing cell type-specific Cre-driver lines with *Trpa1*-floxed mice (*Trpa1^fl/fl^*, exons 22-24 flanked by *loxP* sites) (Fig. 2b). Following Cre-mediated excision of the wild-type Trpa1 exon 22-24, an AAV vector delivers the gRNAs along with a mutated *Trpa1*\* minigene (exons 22-24), allowing targeted re-expression of the modified sequence (*Plp*-Cre-MiniG and *Adv*-Cre-MiniG mice). Expression of SaCas9 in GEdit and MiniG mice was confirmed by immunohistochemistry, detecting GFP fluorescence (Fig. S1 a,b). The presence of the single-point mutation was confirmed by RNA long-reads sequencing in sciatic nerves tissue homogenates, (Fig.S1 c,d).

**Figure 2.**
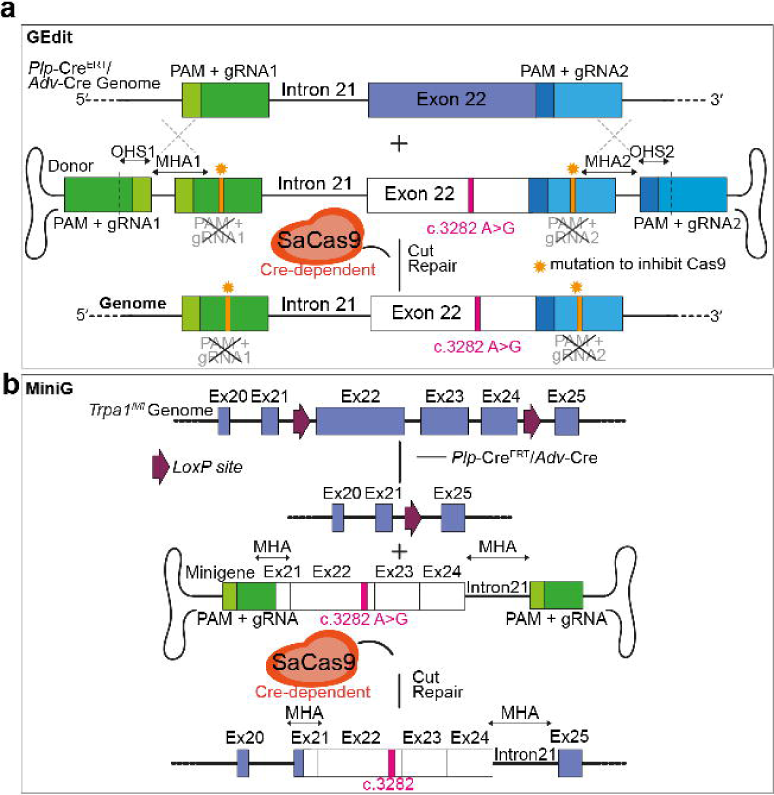
Generation of mouse model with c.3282 A>G mutation on exon 22 of Trpa1 gene. **(a)** Schematic illustration of genome editing approach (GEdit) based on MMEJ mutation substitution. *Plp*-Cre^ERT^ or *Adv*-Cre mice genome and donor DNA with c.3282 A>G mutation on exon 22 carried by AAV were excised at the flanking gRNA target sites (gRNA1 and gRNA2) by SaCas9. Donor DNA with c.3282 A>G is inserted into the mouse genome using microhomology arms (MHA) inducing the FEPS mutation. Single point mutations in gRNA1 and gRNA2 were designed to prevent SaCas9 repeated cleavage after successful replacement. **(b)** Schematic representation of minigene approach (MiniG) based on Cre-*Trpa1^fl/fl^* mice where exons 22-24 of *Trpa1* were flanked by *LoxP* sites. In presence of Cre, exon 22-24 of wild-type *Trpa1* were excised. Minigene with the exons 22*-24 was excised at the flanking gRNA target sites by SaCas9 and inserted into the mouse genome, inducing c.3282 A>G mutation.

### Schwann cell and neuronal-TRPA1* differentially mediate mechanical allodynia and acute nociception

We previously reported that the intraplantar (i.pl.) administration of AITC, the selective TRPA1 agonist, elicited both acute nociception and paw mechanical allodynia, which TRPA1 differentially mediated and expressed in neurons and SCs, respectively^49,50^. To further dissect the contribution of SC and neuronal TRPA1* in the mouse models of FEPS, we employed the i.pl. injection of the two selective TRPA1 agonists, AITC and 4-HNE.

Since patients with FEPS exhibit enhanced sensitization of the nociceptive system upon stimulation^24^, we first identified subthreshold doses of both agonists, that elicited acute nociception and paw mechanical allodynia in FEPS mice but not in Control mice. To this end, Control mice were used to establish a dose-response curve for both AITC and 4-HNE (Fig. 3a,b). We found that administering a subthreshold dose of AITC (0.1 nmol) and 4-HNE (0.5 nmol) induced mechanical allodynia in *Plp*-Cre-GEdit and *Plp-*Cre-MiniG mice, but not in *Adv*-Cre-GEdit, *Adv*-Cre-MiniG, or Control mice (Fig. 3c,d and Fig. S2a,b). This allodynia was abolished by systemic administration of the TRPA1 antagonist A967079 (Fig. 3e and Fig S2c). Conversely, the same subthreshold dose of AITC and 4-HNE evoked an acute nociceptive response in *Adv*-Cre-GEdit and *Adv*-Cre-MiniG mice, but not in *Plp*-Cre-GEdit, *Plp*-Cre-MiniG, or Control mice (Fig 3f,g and Fig. S2d,e). The nociceptive response observed in *Adv*-Cre-GEdit and *Adv*-Cre-MiniG mice was blocked by A967079 (Fig. 3h and Fig. S2f). Collectively, these data reveal that the FEPS-associated TRPA1 mutation in SCs predominantly drives mechanical allodynia. In contrast, the mutation in sensory neurons underlies enhanced acute nociceptive responses.

**Figure 3.**
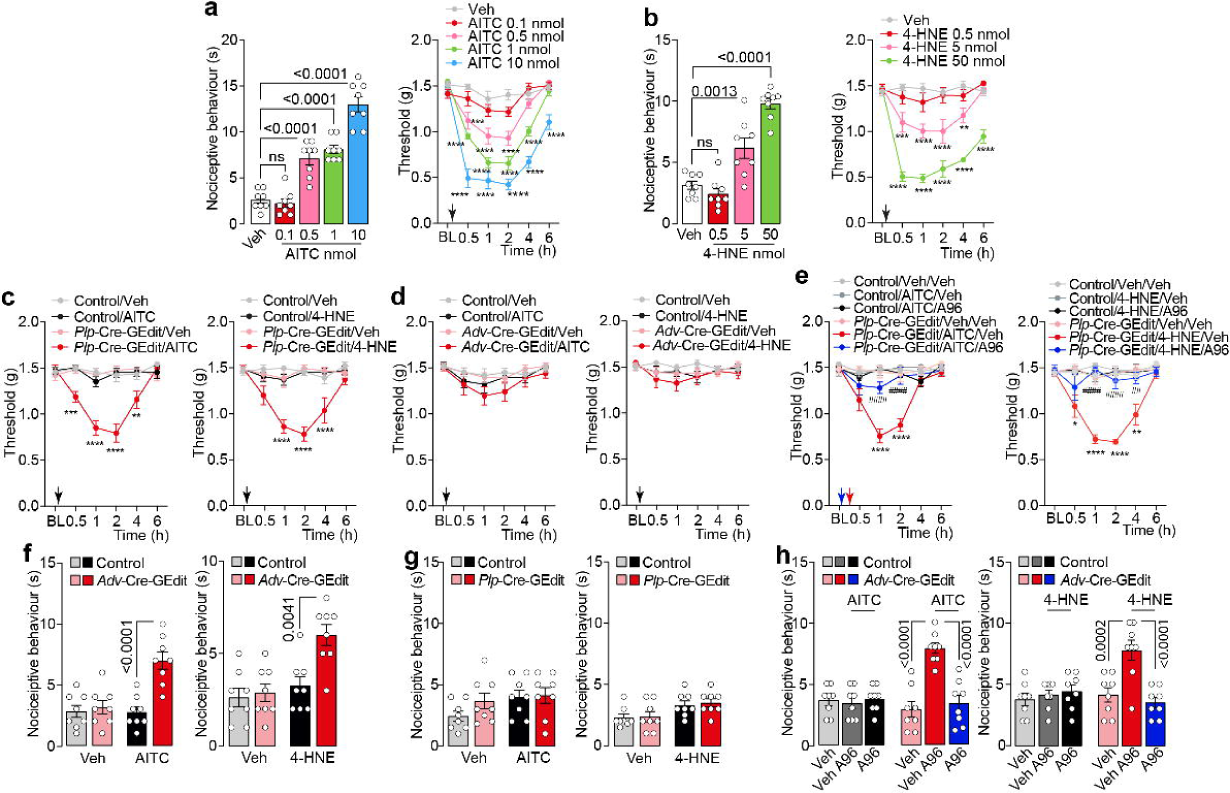
Mutant TRPA1 in Schwann cells and primary sensory neurons independently mediates mechanical allodynia and acute nociception. **(a,b)** Dose-dependent, acute nociception and dose- and time-dependent hind paw mechanical allodynia after intraplantar (i.pl./10 μl) injection of AITC and 4-HNE or vehicle (Veh) in C57BL/6J mice (B6). **(c, d)** Mechanical allodynia after i.pl. AITC (0.1 nmol), 4-HNE (0.5 nmol) or Veh in *Plp-*Cre-GEdit, *Adv-*Cre-GEdit or Control mice. **(e)** Mechanical allodynia after i.pl. AITC (0.1 nmol), 4-HNE (0.5 nmol) or Veh in *Plp-*Cre-GEdit mice pretreated with A967079 (A96, 100 mg/kg, intraperitoneal, i.p.) or Veh. **(f,g)** Acute nociceptive response after i.pl. AITC (0.1 nmol), 4-HNE (0.5 nmol) or Veh in *Adv-*Cre-GEdit, *Plp-*Cre-GEdit or Control mice. **(h)** Acute nociception after i.pl. AITC (0.1 nmol), 4-HNE (0.5 nmol) or Veh in *Adv-*Cre-GEdit mice pretreated with A96 (100 mg/kg, i.p.). Data are mean ± s.e.m. (n=8 mice/group). Arrow indicates time of compounds administration. 1-way and 2-way ANOVA, Bonferroni correction. *p<0.05, **p<0.01, ***p<0.001, ****p<0.0001 vs Veh or vs Control/Veh or vs Control/Veh/Veh; ^##^p<0.01, ^####^p<0.0001vs *Plp-*Cre-GEdit/HNE/AITC/Veh.

### Conditional knock-in model confirms dichotomous roles of SC and neuronal TRPA1*

To further validate the distinct roles of SCs and sensory neurons in FEPS, we designed a conditional knock-in (cKI) mouse model expressing the heterozygous FEPS mutation. The cKI-TRPA1 mouse model was created using an advanced Cre-dependent genetic switch strategy^51^. This Cre-*loxP* technology-based method enables a shift in genetic expression from the wild-type mouse TRPA1 to the mutated TRPA1*. To generate the cKI-TRPA1* mouse, we constructed a plasmid to electroporate C57BL/6N embryonic stem (ES) cells, containing *loxP*-flanked DNA sequence from mouse exon 22 in the sense orientation and the mutated and degenerated exon 22 in the antisense orientation (Fig 4a and Fig. S3a-h). Following selective Cre recombination, the mouse wild-type exon was excised, and the mutated exon was inverted to produce *Plp*-cKI-*Trpa1\** and *Adv*-cKI-*Trpa1\** mice that harbor mutant *Trpa1\** exon specifically in SCs and sensory neurons, respectively (Fig 4a).

**Figure 4.**
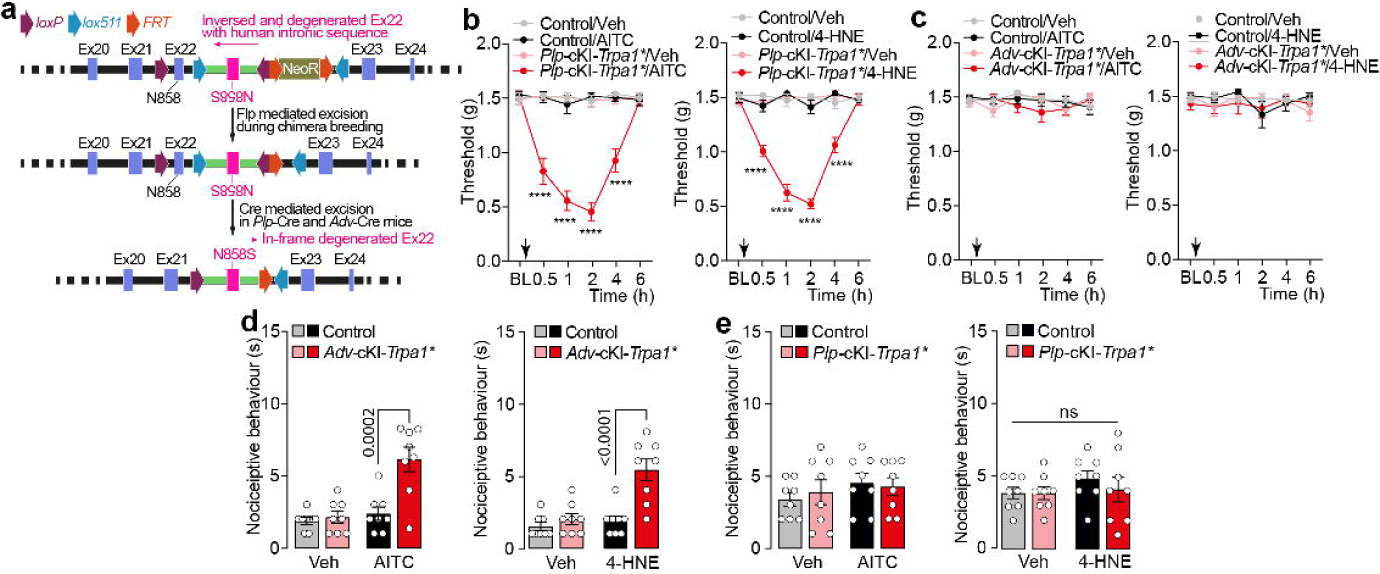
Knock-in FEPS mouse model reveals specific role of mutant TRPA1 in Schwann cells and primary sensory neurons in mechanical allodynia and acute nociception. **(a)** Schematic representation of conditional Knock-in FEPS mouse model. **(b,c)** Time-dependent mechanical allodynia after intraplantar (i.pl./10 μl) injection of AITC (0.1 nmol), 4-HNE (0.5 nmol) or vehicle (Veh) in *Plp*-cKI-*Trpa1*, Adv*-cKI-*Trpa1\** or Control mice. **(d,e)** Acute nociception after i.pl. AITC (0.1 nmol), 4-HNE (0.5 nmol) or Veh in *Adv*-cKI-*Trpa1*, Plp*-cKI-*Trpa1\** or Control mice. Data are mean ± s.e.m. (n=8 mice/group). Arrow indicates time of compounds administration. 2-way ANOVA, and 1-way ANOVA, Bonferroni correction. ****p<0.0001 vs Control/Veh

Subthreshold dose (i.pl.) of AITC and 4-HNE triggered mechanical allodynia in *Plp*-cKI-*Trpa1*\* mice but not in *Adv*-cKI-*Trpa1*\* or Control mice (Fig 4b,c). Conversely, the same agonists induced nociceptive behaviour in *Adv*-cKI-*Trpa1\** mice but not in *Plp*-cKI-*Trpa1\** or Control mice (Fig. 4d,e). The point mutation harboured by Plp-cKI-*Trpa1\** and *Adv*-cKI-*Trpa1\** mice was confirmed by next-generation sequencing (Fig. S3i).

These findings confirm the dichotomous roles of SCs and neuronal TRPA1 in mediating mechanical allodynia and nociception, respectively. Importantly, the study established novel transgenic mouse models to investigate FEPS and TRPA1-related pain syndromes.

### Fasting, cold and stress exposure trigger Schwann cell TRPA1-dependent allodynia in FEPS mice

Individuals with FEPS experience recurrent episodes of severe upper body pain beginning in infancy and persist throughout life. These painful crises are notably precipitated by physiological or environmental stressors such as prolonged food restriction, emotional or physical stress, or exposure to low temperatures^24^. To investigate the mechanisms and specific cells involved in pain episodes in FEPS, we tested the mechanical allodynia in transgenic mice under physiological and environmental stressors, including fasting, cold exposure, and restraint stress^52–55^.

Firstly, we found that neither 24 nor 48 hours of food deprivation induced mechanical allodynia in Control mice (Fig. 5a,b). In contrast, *Plp*-Cre-GEdit, *Plp*-Cre-MiniG and *Plp*-cKI-*Trpa1\** mice developed mechanical allodynia beginning at 24 hours of food deprivation (Fig. S4a and Fig. 5c). Conversely, *Adv*-Cre-GEdit, *Adv*-Cre-MiniG and *Adv*-cKI-*Trpa1\** mice did not exhibit paw mechanical allodynia following food deprivation (Fig. S4b and Fig. 5d). We next found that the exposure to cold temperature (4 °C) for 15 hours did not induce mechanical allodynia in Control mice when assessed up to 4 hours after the exposure (Fig. 5e,f). In contrast, cold temperature, induced time-dependent mechanical allodynia in *Plp*-Cre-GEdit, *Plp*-Cre-MiniG and *Plp*-cKI-*Trpa1\** mice but not in Adv-Cre-GEdit, Adv-Cre-MiniG or Adv-cKI-*Trpa1\** mice (Fig. S4c,d and Fig. 5g,h). Finally, following a single 2-hours restraint stress, *Plp* -Cre-GEdit, *Plp*-Cre-MiniG mice and *Plp*-cKI-*Trpa1\**, but not *Adv*-Cre-GEdit, *Adv*-Cre-MiniG, *Adv*-cKI-*Trpa1\** and Control mice, developed mechanical allodynia up to 24 hours post-restraint (Fig. S4e,f and Fig. 5i-l). Together, these findings underscore the role of SC TRPA1*\** as a key modulator of mechanical allodynia triggered by fasting, cold exposure, and restraint stress in the mouse model of FEPS.

**Figure 5.**
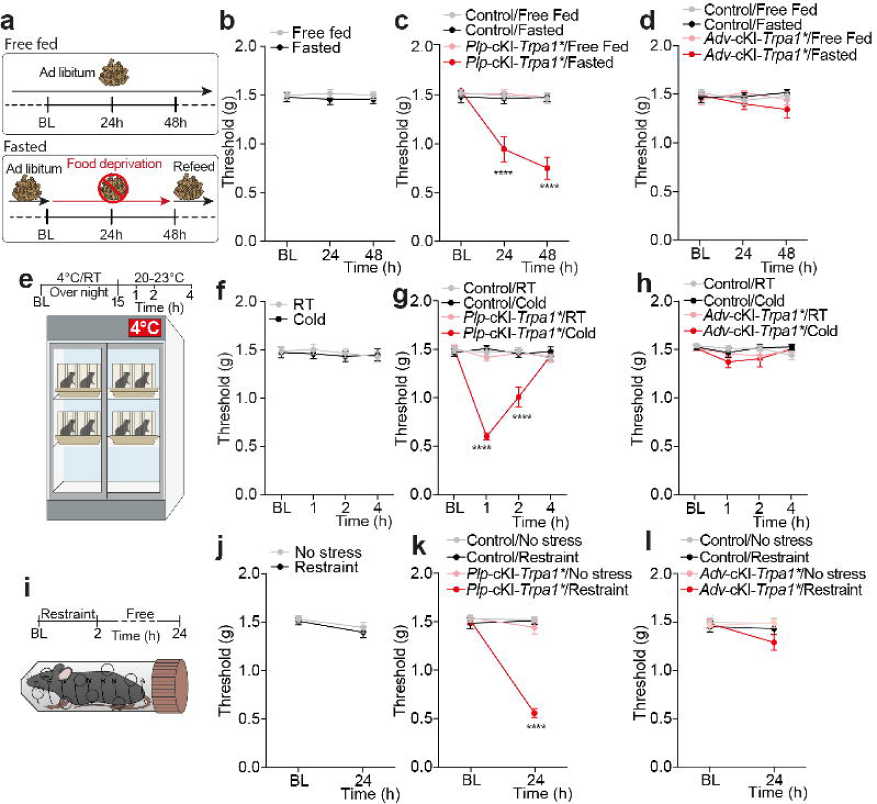
Fasting, cold exposure, and restraint stress induce Schwann cell-dependent mechanical allodynia mediated by mutant TRPA1. **(a)** Schematic representation of free or fasted diet regimen. **(b-d)** Time dependent mechanical allodynia in free fed or fasted *Plp*-cKI-*Trpa1\**, *Adv*-cKI-*Trpa1\** or Control mice. **(e)** Schematic representation of cold (4 °C) or room temperature (RT) exposure **(f-h)** Time dependent mechanical allodynia in *Plp*-cKI-*Trpa1\**, *Adv*-cKI-*Trpa1\** or Control mice (n=8 mice/group) exposed to cold or RT. **(i)** Schematic representation of restraint stress model. **(j-l)** Time dependent mechanical allodynia in the presence or absence of restraint stress in *Plp*-cKI-*Trpa1\**, *Adv*-cKI-*Trpa1\** or Control mice. Data are mean ± s.e.m. (n=8 mice/group). 2-way ANOVA, Bonferroni correction. ****p<0.0001 vs Control/Fasted, Control/Cold, Control/Restraint.

### Schwann cell TRPA1* amplifies oxidative stress to sustain mechanical allodynia

The development and maintenance of chronic pain have been linked to oxidative stress/ROS levels within the peripheral nervous system across several animal models of pain^56–59^. We previously reported that SC TRPA1 drives feed-forward mechanism that amplifies and sustains oxidative stress-related nociceptive signalling, thereby contributing to the persistence of mechanical allodynia^45^). Having disentangled the distinct contribution of neuronal and SC TRPA1 to acute nociception and sustained mechanical allodynia, respectively, we investigated the role of oxidative stress in the development of mechanical allodynia in FEPS mice.

Firstly, we assessed ROS production in TRPA1 and TRPA1*-HEK293T cells following stimulation with hydrogen peroxide (H_2_O_2_), AITC or 4-HNE. In both TRPA1 or TRPA1*-HEK293T transfected cells, H_2_O_2_, AITC and 4-HNE elicited a concentration-dependent increase in intracellular H_2_O_2_ levels with significative higher potency and efficacy on TRPA1*-HEK293T. These effects were reduced in the presence of A967079 (30 μM) (Fig. 6a-c).

**Figure 6.**
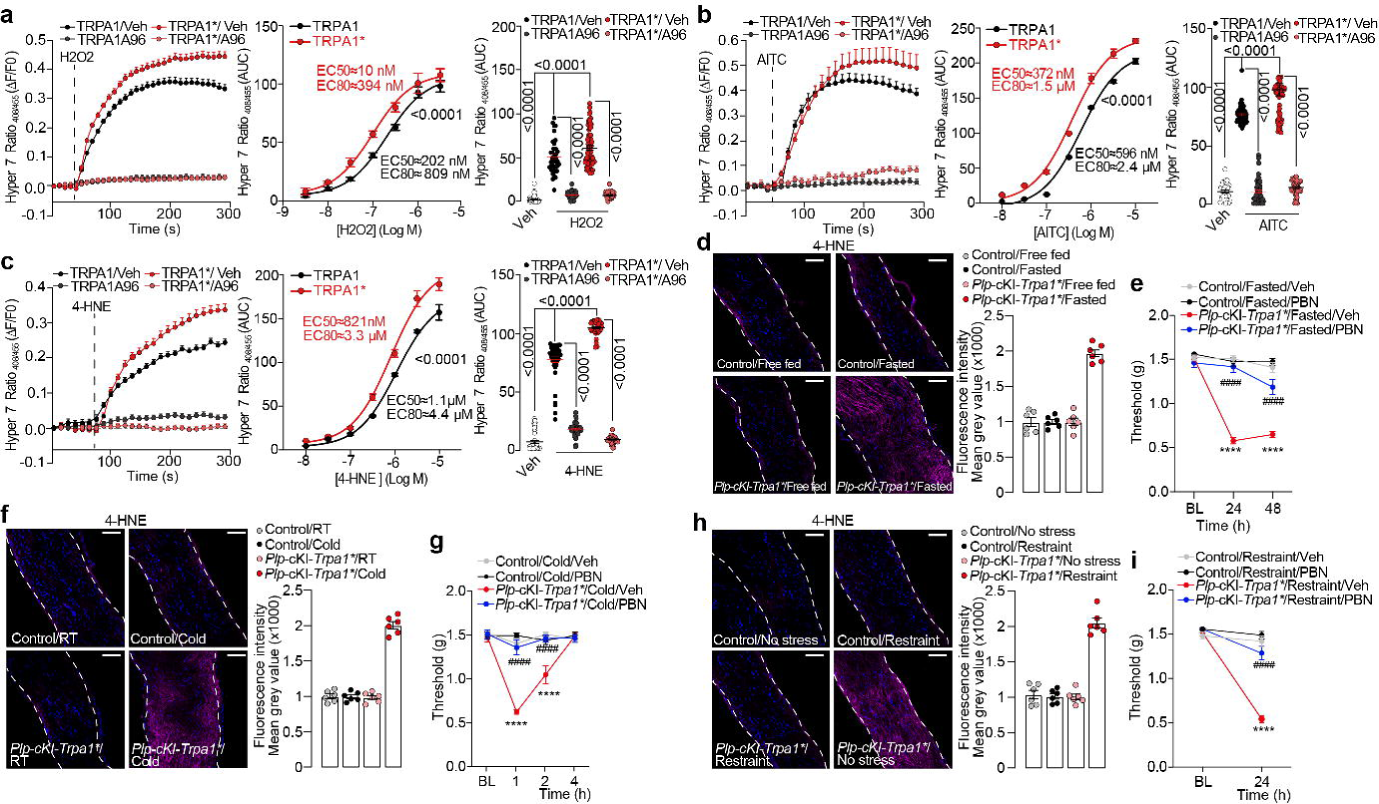
Mutant TRPA1* channel generates oxidative stress to maintain mechanical allodynia. **(a,b,c)** Typical traces, concentration response curve (CRC) and pooled data of HyPer7.2 in HEK293T cells transfected with the mouse TRPA1 channel (TRPA1) or its mutated counterpart (TRPA1*), stimulated with hydrogen peroxide (H_2_O_2_, 3 nM-3 µM), AITC (10 nM-10 µM) and 4-HNE (10 nM-10 µM) also in the presence of A967079 (A96, 30 µM) or vehicle (Veh, 0.0001% DMSO) (cells number: H_2_O_2_ CRC TRPA1: 3 nM n= 56, 10 nM n=46, 30 nM n=57, 100 nM n=66, 300 nM n=61, 1 µM n=66, 3 µM n=65. EC_50_CI=149-275 nM, EC_80_CI=598 nM-1.1 μM. H_2_O_2_ CRC TRPA1*: 3 nM n= 50, 10 nM n=103, 30 nM n=66, 100 nM n=63, 300 nM n=43, 1 µM n=47, 3 µM n=56. EC_50_CI=7-144 nM, EC_80_CI=271-577 nM. H_2_O_2_ cumulative data: Veh n=48, TRPA1/Veh n=42, TRPA1/A96 n=66, TRPA1*/Veh n=67, TRPA1*/A96 n=66; AITC CRC TRPA1: 10 nM n= 93, 30 nM n=87, 100 nM n=55, 300 nM n=73, 1 µM n=68, 3 µM n=55, 10 µM n=88. EC_50_CI=549-649 nM, EC_80_CI=2.2-2.6 μM. AITC CRC TRPA1*: 10 nM n=36, 30 nM n=31, 100 nM n=43, 300 nM n=71, 1 µM n=66, 3 µM n=55, 10 µM n=57. EC_50_CI=311-444 nM, EC_80_CI=1.2-1.8 μM. AITC cumulative data: Veh n=31, TRPA1/Veh n=73, TRPA1/A96 n=64, TRPA1*/Veh n=61, TRPA1*/A96 n=50; 4-HNE CRC TRPA1: 10 nM n= 55, 30 nM n=60, 100 nM n=47, 300 nM n=56, 1 µM n=40, 3 µM n=46, 10 µM n=37. EC_50_CI=926 nM-1.3 μM, EC_80_CI=3.7-5.3 nM. 4-HNE CRC TRPA1*: 10 nM n=82, 30 nM n=46, 100 nM n=50, 300 nM n=35, 1 µM n=23, 3 µM n=61, 10 µM n=66. EC_50_CI=617 nM-1.2 μM, EC_80_CI=2.5-4.4 μM. 4-HNE cumulative data: Veh n=39, TRPA1/Veh n=44, TRPA1/A96 n=48, TRPA1*/Veh n=38, TRPA1*/A96 n=46). **(d,f,h)** Representative images and cumulative data of 4-HNE content within the sciatic nerve of *Plp*-cKI-*Trpa1\**or Control mice **(d)** free fed or fasted, **(f)** exposed to cold or room temperature (RT) or **(h)** in the presence or absence of restraint stress. (n=6, scale bar: 50 μm). **(e,g,i)** Time dependent mechanical allodynia following repeated administration of phenyl-N-tert-butylnitrone (PBN, 100 mg/kg,intraperitoneal, i.p.) or vehicle (Veh) in *Plp*-cKI-*Trpa1\** or Control mice **(e)** fasted, **(g)** exposed to cold or **(i)** restraint stress. Data are mean±s.e.m. (n=8 mice/group) 1-way and 2-way ANOVA, Bonferroni correction. ****p<0.0001 vs Control/Fasted/Veh, Control/Cold/Veh, Control/Restraint/Veh; ####p<0.0001vs *Plp*-cKI-*Trpa1\**/Fasted/Veh, *Plp*-cKI-*Trpa1\**/Cold/Veh, *Plp*-cKI-*Trpa1\** /Restraint/Veh

Fasting, cold exposure and restraint stress conditions evoked mechanical allodynia in *Plp*-Cre-GEdit, *Plp*-Cre-MiniG and *Plp*-cKI-*Trpa1\** but not in *Adv*-Cre-GEdit, *Adv*-Cre-MiniG, *Adv*-cKI-*Trpa1\** and Control mice. The same conditions also induced accumulation of 4-HNE within the sciatic nerve of mice carrying the *Trpa1\** mutation in SCs cells, but not in those with a mutation restricted to sensory neurons (Fig. 6d,f,h and Fig. S4g-i). To confirm the role of oxidative stress in FEPS mice, we also tested the effect of the antioxidant phenyl-N-tert-butylnitrone (PBN). Repeated administration of PBN reversed mechanical allodynia in *Plp*-Cre-GEdit, *Plp*-Cre-MiniG and *Plp*-cKI-*Trpa1\** mice following fasting, cold exposure, or restraint stress (Fig. 6e,g,i and Fig. S4j-l). We speculate that the mutated TRPA1 in Schwann cells detected subthreshold levels of oxidative stress in FEPS mice subjected to different stress conditions. Subsequent activation of TRPA1*\** in Schwann cells triggered a feed-forward mechanism that amplifies and sustains mechanical allodynia.

## Discussion

Here, we provide evidence that, in our mouse models of FEPS, the gain-of-function mutation of the TRPA1 channel exerts a strikingly cell-specific role in pain modulation. By studying TRPA1* in Schwann cells and neurons, we demonstrate that the mutation differentially regulates acute nociception and chronic pain states. Specifically, TRPA1* expression in primary sensory neurons enhances the perception of acute painful stimuli, whereas its expression in Schwann cells drives the development and persistence of mechanical allodynia. These findings uncover an unexpected functional dichotomy in TRPA1 signaling and identify Schwann cells as a key cellular substrate through which TRPA1 gain-of-function mutations can promote chronic pain.

Pain research has traditionally focused on neurons as the primary mediators of nociceptive transmission. However, increasing evidence indicates the critical role of non-neuronal cells in the initiation and maintenance of pain states. Our findings do not contradict the dogma that neurons are the final effectors of pain signals; rather, upon injury or insult, non-neuronal cells release neuromodulatory substances in proximity to nociceptors, thereby sustaining pain^60^. Following nerve injury, immune cells can infiltrate the peripheral nervous system, releasing pro-algesic factors such as leukocyte elastase from T lymphocytes or tumor necrosis factor, interleukin-1β, and ROS from monocytes/macrophages^61–63^. In parallel, non-immune resident cells such as keratinocytes and satellite glial cells secrete neuroactive molecules, such as prostaglandin E2, ATP, nerve growth factor, and matrix metalloproteinases, that directly modulate nociceptor excitability^64–67^.

We have previously shown that Schwann cells, the primary glial cells of the peripheral nervous system, acquire a pro-algesic phenotype in multiple mouse models of pain, releasing ROS that sensitize nearby nociceptors.^36,37,39,41,42,45,68^. In this study, we extend the role of Schwann cells in a genetically driven pain syndrome, demonstrating that TRPA1 gain-of-function signaling in these cells is sufficient to maintain persistent mechanical allodynia independently of neuronal TRPA1* expression. To dissect the cell-specific contribution of TRPA1* in FEPS, we generated three complementary transgenic mouse models of FEPS carrying the TRPA1* mutation in Schwann cells and primary sensory neurons. We found that i.pl. administration of subthreshold doses of two different selective TRPA1 agonists (AITC and 4-HNE) elicited a robust acute nociceptive response in mice expressing the TRPA1* in primary sensory neurons that was ineffective in wild-type mice. In contrast, mice expressing TRPA1* in Schwann cells developed pronounced mechanical allodynia, a phenotype absent in both wild-type mice and in animals expressing TRPA1* in primary sensory neurons.

Individuals suffering from chronic pain frequently avoid conditions or activities that could worsen their symptoms. It is widely reported that prolonged or excessive stress, including sleep disturbances, can induce dysfunction of cortisol activity, leading to widespread inflammation and enhanced pain sensitivity^69,70^. Similarly, barometric changes trigger imbalances in brain chemicals, including serotonin, which facilitate migraine attacks^71^. Acute physical activity and exercise induce phosphorylation of the NMDA receptor in peripheral nerves, a mechanism that can promote pain sensitization in chronic pain conditions^72^. Importantly, patients with familial FEPS report that episodes of severe pain are often precipitated by physiological or environmental stressors, including fasting, cold exposure, or psychological stress^24^. In line with these observations, we modeled food deprivation, cold exposure, and restraint-induced stress in mice. Under these subthreshold discomfort conditions, mice expressing TRPA1* selectively in primary sensory neurons, did not develop mechanical allodynia, similarly to wild-type controls. In sharp contrast, mice expressing TRPA1* in Schwann cells displayed robust pain behaviors under the same conditions, indicating that Schwann cell-mediated TRPA1 signaling is sufficient to convert mild environmental or metabolic stressors into prolonged painful responses, highlighting a critical role for non-neuronal TRPA1 activity in FEPS pathogenesis. The contribution of ROS in chronic pain is widely recognized^73–77^. We have previously demonstrated that Schwann cells TRPA1 plays a critical role in ROS production and amplification, a process tightly correlated to the onset and persistence of chronic pain. Several pro-algesic mediators, such as CGRP in periorbital allodynia, CSF-1 in primary tumor, IGF-1 in bone metastatic pain, and C5a in endometriosis pain, activate intracellular pathways in Schwann cells that ultimately lead to TRPA1 activation, excessive ROS generation, and paracrine sensitization of primary sensory neurons^37,41,46^. Here, we show that the TRPA1* channel was activated by subthreshold doses of ROS, leading to TRPA1-dependent H_2_O_2_ release, an effect not observed in wild-type TRPA1. Moreover, *in vivo* models of food deprivation, cold exposure, and constraint-induced stress resulted in ROS accumulation in the peripheral nerves of mice with Schwann cell TRPA1*, but not in wild-type mice or in animals carrying the TRPA1 mutation in primary sensory neurons. We speculate that these mild stress conditions activated intracellular pathways in Schwann cells that remain below the activation threshold of wild-type TRPA1 but are sufficient to engage TRPA1*, thereby initiating a feed-forward mechanism of ROS production and sustained nociceptor sensitization.

In conclusion, our study dissects the contributions of TRPA1 in Schwann cells and primary sensory neurons to mechanical allodynia and acute pain in FEPS mice. It provides complementary transgenic models to investigate the distinct roles of neuronal and non-neuronal TRPA1 signaling in chronic pain disorders.

### Limitations of the Study

While this study provides new insights into the cellular mechanisms underlying pain in FEPS patients, it does not directly offer a therapeutic strategy capable of providing long-term relief from chronic pain. Nonetheless, the animal model established here represents a valuable platform for the development of innovative biotechnological approaches, such as selective silencing of TRPA1 expression in Schwann cells or gene therapy strategies to modify the genetic sequence, aimed at reducing cellular hypersensitivity to discomfort in FEPS patients.

## RESOURCE AVAILABILITY

### Lead contact

Further information and requests for resources and reagents should be directed to and will be fulfilled by the lead contacts Romina Nassini; Francesco De Logu

### Materials availability

All materials used in this study are available to any researcher for purposes of reproducing or extending the analyses. This study did not generate any new, unique reagents.

### Data and code availability

Data Availability: Fastq sequencing data generated in this study are available in the Sequence Read Archive (SRA) under BioProject accession number PRJNA1424206.

Code Availability: No custom code was used within the study.

## ACKNOWLEDGMENTS

This work was sponsored by Fondazione Telethon (Grant no GMR22T1070) (FDL)

## AUTHOR CONTRIBUTIONS

Conceptualization, EC, MMo, KYK, MCB, RL, RN, FDL; data curation, MM, MC, EC, LB, LL, IS, RN, FDL; formal analysis, MM, MC, EC, AP, GDS, EB, LT, VA, GF; funding acquisition, FDL; investigation, MM, MC, EC, MMo, LB, LL, IS, AP, GDS, EB, LT, VA, GF, EDdoNM, MCB, RL, RN, FDL; methodology, MM, MC, EC, MMo, GDS, MCB, RL; project administration, RN, FDL; resources, RN, FDL; supervision; RN, FDL; writing – original draft, MM, MC, EC, MMo, KYK, MCB, RL, RN, FDL; writing – review & editing, MM, MC, EC, KYK, MCB, RN, FDL

## Methods

### Animals

Male mice were used throughout (25-30 g, 7-9 weeks). In accordance with the 3R guidelines to minimize the number of animals and avoid possible confounding effects of hormone fluctuation in pain perception, only male mice were used. The following mouse strain was used: C57BL/6J mice (Charles River, RRID: IMSR_JAX:000664).

The C57BL/6N-*Trpa1^tm1.1Ics^* (*Trp1a^cKI^*) mutant mouse line was generated at the Institut Clinique de la Souris - PHENOMIN (http://www.phenomin.fr). To produce the conditional N855S knock-in Trpa1 mouse model, a targeting plasmid was engineered and electroporated into C57BL/6N embryonic stem (ES) cells. The construct contained two distinct pairs of recombination sites, one pair of *wild-type loxP* sites and one pair of *lox511* sites, each arranged in a head-to-head orientation. The targeting vector included the murine exon 22 sequence together with its flanking intronic regions cloned in the sense orientation, and a modified “degenerated” human exon 22 sequence harboring the N855S mutation along with its corresponding human intronic flanking regions cloned in the antisense orientation (**Fig. 4a**). Both *loxP* and *lox511* sites were positioned in an inverted orientation and are recognized by Cre recombinase; however, *lox511* sites only recombine efficiently with identical *lox511* sites and not with *loxP* sites. Upon Cre-mediated recombination, an initial inversion event occurs at either the *loxP* or *lox511* sites, which are present in opposite orientations, resulting in the formation of two identical recombination sites (either *loxP*–*loxP* or *lox511*–*lox511*) in the same orientation. Subsequent Cre activity leads to excision of the intervening DNA sequence between the recombination sites. This recombination strategy ultimately yields an allele containing a single *loxP* site and a single *lox511* site, thereby preventing further inversion events. Following recombination, the endogenous promoter drives expression of the mutant *Trpa1* allele in place of the *wild-type Trpa1* gene. To introduce *Trpa1* N855S point mutation in Schwann cells or primary sensory neurons, homozygous *Trpa1^cKI^* were crossed with hemizygous *Plp-Cre^ERT^* mice or hemizygous *Adv-Cre* mice, respectively, as for the *Trpa1^fl/fl^* mice, homozygous *Trpa1^cKI/cKI^*mice (*Plp-Cre^ERT^; Trpa1^cKI/cKI^ and Plp-Cre^ERT–^; Trpa1^cKI/cKI^*, respectively) were subjected to intraperitoneal (i.p.) injection of 4-hydroxytamoxifen (4-OHT, 1□mg/100□μL in corn oil once a day consecutively for 3 days)^78^. To generate mice in which the *Trpa1* gene was conditionally silenced in Schwann cells, homozygous 129S-Trpa1^tm2Kykw/J^ (floxed Trpa1, *Trpa1^fl/fl^*, RRID:IMSR_JAX: 008649 Jackson Laboratory) were crossed with hemizygous B6.Cg-Tg(Plp1-CreERT)3Pop/J mice (*Plp-Cre^ERT^*, RRID: IMSR_JAX:005975 Jackson Laboratory) expressing a tamoxifen-inducible Cre in Schwann cells (Plp1, proteolipid protein myelin 1). The progeny (*Plp-Cre^ERT^;Trpa1^fl/fl^*) was genotyped using PCR for *Trpa1* and *Plp-Cre^ERT^.* Mice that were negative for *Plp-Cre^ERT^* (*Plp-Cre^ERT–^;Trpa1^fl/fl^*) were used as control. Both positive and negative mice for *Cre^ERT^* and homozygous floxed Trpa1 (*Plp-Trpa1* and control respectively) were treated with 4-OHT, 1□mg/100□μL, i.p., once a day consecutively for 3 days. To selectively delete *Trpa1* in primary sensory neurons, Trpa1^fl/fl^ mice were crossed with hemizygous Advillin-Cre mice (Adv-Cre) ^79^. Mice positive or negative for Cre and homozygous for floxed Trpa1 (*Adv-Trpa1;Trpa1^fl/fl^* and control respectively) were used. The successful Cre-driven deletion of TRPA1 mRNA was confirmed using reverse transcription quantitative real-time PCR (RT-qPCR).

*Plp-Cre^ERT^, Adv-Cre, Plp-Cre^ERT^;Trpa1^fl/fl^, Adv-Trpa1;Trpa1^fl/fl^* and control mice were infected with an intravenous or intrathecal (i.v. or i.th., 10 µL, 1×10^12^ v/g) injection of different AAVs and animals were used 3 weeks after AAVs infection.

The group size of n=8 animals for behavioral experiments was determined by sample size estimation using G*Power (v3.1;^80^) to detect size effect in a post-hoc test with type 1 and 2 error rates of 5 and 20%, respectively. Mice were allocated to vehicle or treatment groups using a randomization procedure (http://www.randomizer.org/). The assessors were blinded to the identity of the animals (allocation to treatment groups). None of the animals were excluded from the study. Behavioral studies followed Animal Research: Reporting of In Vivo Experiments (ARRIVE)^81^. Animals were euthanized with inhaled CO_2_ plus 10-50% O_2_. Mice were housed in a temperature (20 ± 2 °C) and humidity (50 ± 10%) controlled *vivarium* (12 h dark/light cycle, free access to food and water, five animals per cage). At least 1 h before behavioral experiments, mice were acclimatized to the experimental room and behavior was evaluated between 9:00 am and 5:00 pm. All the procedures were conducted following the current guidelines for laboratory animal care and the ethical guidelines for investigations of experimental pain in conscious animals set by the International Association for the Study of Pain^82^.

### Treatment Protocols

If not otherwise indicated, the reagents were obtained from Merck Life Science SRL (Milan, Italy). C57BL/6J mice received intraplantar (i.pl./10 µl site) injection of allyl isothiocyanate (AITC, 0.1, 0.5, 1, 10 nmol) or 4-hydroxynonenal (4-HNE, 0.5, 5, 50 nmol) or their vehicle (1% dimethyl sulfoxide, DMSO, in 0.9% NaCl); A967079 (100 mg/kg) or its vehicle (1% DMSO in 0.9% NaCl) were administered intraperitoneal (i.p., 10 ml/kg). α-Phenyl-N-tert-butylnitrone (PBN, 100 mg/kg, i.p.) or its vehicle (2% DMSO in 0.9% NaCl) were administered before and subsequently once daily for two consecutive days in mice subjected to fasting, cold, and stress exposure.

### Behavioural experiments

#### Acute nociception

Immediately after i.pl. injection, mice were placed inside a plexiglass chamber, and spontaneous nociception was assessed for 10□min by measuring the time (seconds) that the animal spent licking/lifting the injected paw.

#### Hindpaw mechanical allodynia

The mechanical paw-withdrawal threshold was measured using von Frey filaments of increasing stiffness (0.02-2 g) applied to the plantar surface of the mouse hind paw, according to the up-and-down paradigm^83^. The 50% mechanical paw-withdrawal threshold (g) response was then calculated from the resulting scores.

#### Fasting schedule

After baseline mechanical threshold assessment, mice were deprived of the standard pellet diet for 48 h. Water was available *ad libitum* throughout the entire experimental period. Behavioral testing was performed every 24 h during the fasting period and after food deprivation.

#### Single Restraint

Mice were placed in cylindrical tail-access rodent restrainers designed for animals weighing 15-30 g (Stoelting, 51338). Animals were positioned in the restraint devices to prevent movement without impairing normal respiration. Restraint was maintained for 2 h following baseline mechanical threshold assessment. Behavioral testing was conducted 24 h after the restraint time. Control mice were left in their home cages for 2 h without access to food or water to control for the effects of food and water deprivation alone on stress-related response^84^.

#### Cold exposure

Mice were housed in plastic cages with *ad libitum* access to food and water and placed in a temperature-controlled environment at 4 ± 2 °C for 14-15 h (overnight). Prior to behavioral testing, all mice were transferred to the experimental room (20-23 °C) and allowed to acclimate for at least 30 min, after which behavioral experiments were performed^55^.

### Cell line

#### HEK293T cell line

Human kidney epithelial (HEK293T) cells (HEK293T; #CRL3216™; ATCC, Manassas, VA, USA RRID:CVCL0063) were cultured in DMEM supplemented with fetal bovine serum (FBS, 10%) and L-glutamine (2 mM) at 37 °C in 5% CO_2_ and 95% O_2_.

#### AAVpro 293T cells

AAVpro 293T cells (#632273, Takara), were maintained in Dulbecco’s Modified Eagle Medium (DMEM) high glucose supplemented with heat inactivated FBS (10%), sodium pyruvate (1 mM), L-glutamine (4 mM) and penicillin/streptomycin (1 mM) in 5% CO_2_ and 95% O_2_ at 37°C. The day before transfection, cells were plated in DMEM supplemented with FBS (2%).

### Plasmids

Plasmids expressing TRPA1 and TRPA1* for in vitro experiments were purchased from VectorBuilder (pRP CMV 3xFLAG-mTrpa1-T2A-Puro VB230123-1262qaf and pRP CMV 3XFLAG-mTRPA1 MUT-T2a-Puro VB230412-1022ruq). To generate mice harbouring mutated copies of the TRPA1 channel, a genome-editing (GEdit) approach and a minigene (MiniG) approach were used. For the GEdit strategy, the plasmid pAAV[Exp]-{Donor_2xsgRNA_TRPA1}-CMV>mCherry (VB230506-1032wnn) was utilized. For the MiniG strategy, the plasmid pAAV[Exp]-{miniGene-DonorMMEJ_gRNA-hTRPA1}-CMV>mCherry (VB230506-1057nhs) was employed. Both approaches were carried out in conjunction with the SaCas9 expression plasmid pAAV[Exp]-{CMV short form}>LL/rev(SaCas9(ns)/T2A/EGFP)/rev(LL) (VB230508-1094rfa). All plasmids were obtained from VectorBuilder.

### AAV Generation

#### AAV production and cell lysis

Three days before Transfection, AAVpro293T cells were seeded in a CellBIND Polystyrene CellSTACK 2 Chamber (#3310, Corning, Corning, NY, USA) for 48 h to reach 80% confluence. The day before transfection, 250 million cells were seeded in a CellBIND Polystyrene CellSTACK 5 Chamber (#3311; Corning) for 24 h, until 80% of confluence. AAVpro 293T cells were triple transfected with polyethylenimine (#23966, PEI, Polyscience) using a DNA:PEI ratio of 1:3. transfection was performed with 2.5 mg total DNA with 1:1:1 molar ratio of three plasmids (plasmid expressing gene of interest, pAdDeltaF6; (#11287, Addgene) and Rep/Cap plasmid). To infect sensory neurons and Schwann cells with high efficiency, the pAAV2/9n and AAV2/rh10 plasmids were used, respectively (pAAV2/9n, #112865; pAAV Rep2/Cap-rh10, #112866, Addgene). After 72 h, AAV virions were collected. Cells were harvested, transferred to 50 mL conical tubes, and centrifuged at 1200 × g for 10 min at 4 °C. The supernatant was filtered through 0.45 μm PES membrane. To concentrate the virus, 25 mL of 40% PEG solution (pH 7.4; 400 g of PEG 8000 + 24 g of NaCl in ddH_2_O to a final volume of 1 mL) was added to every 100 mL of filtered supernatant. The solution was slowly stirred at 4 °C for 1 h and then kept for 3 hrs without stirring at 4 °C to allow complete precipitation of AAV particles. After centrifugation at 2900 × g for 15 min at 4 °C, the supernatant was discarded, and the pellet containing virion particles was resuspended in 10 ml of PBS/pluronic F68 (0.001%)/NaCl (200 mM). AAVpro293T cell pellet, derived from the first centrifugation, was directly resuspended in 10 mL of PBS/Pluronic F68 (0.001%)/NaCl (200 mM) solution and lysed by 4 cycles of freezing at −80°C for 30min, followed by a thaw at 37°C for 10 min. After centrifugation at 3.200 × g for 15 min at 4 °C, supernatant containing the AAV particles was collected. Both samples, obtained from the supernatant and cell pellet, were mixed and incubated with Benzonase (50 U/ml) at 37°C for 45 min to remove residual DNA carried over from the AAV packaging process. The sample was then centrifuged at 2.500 × g for 10 min at 4°C, and the clarified supernatant was transferred to a new tube and kept overnight at 4°C before iodixanol purification.

#### AAV purification by Iodixanol gradient

An iodixanol gradient was prepared. Starting with a 60% iodixanol solution (OptiPrep; STEMCELL Technologies, Vancouver, Canada), 15% iodixanol solution [4.5 mL of iodixanol (60%) + 13.5 mL of NaCl/PBS-MK buffer (1M)], a 25% solution [5 mL of iodixanol (60%) + 7 mL of PBS-MK buffer (1×) + 30 μL of phenol red], a 40% solution [6.7 mL of iodixanol (60%) + 3.3 mL of PBS-MK buffer (1×)], and a 60% solution [10 mL of iodixanol (60%) + 45 μL of phenol red] were made. Each solution was added into a 39-mL Quick-Seal tube (Beckman Coulter, Brea, CA, USA) using an 18 G needle syringe in the following order: 8 mL of 15% iodixanol solution, 6 mL of 25% iodixanol solution, 5 mL of 40% iodixanol solution, and 5 mL of 60% iodixanol solution. Finally, Quick-Seals were filled with AAV solution from the top, sealed, and ultracentrifuged in a Type 70 Ti rotor (Beckman Coulter) at 350.000 x g at 10°C for 1.5 h. Then, Quick-Seal tubes were pierced with a 16 G needle on top and a 14 G needle at the interface between the 60% and 40% iodixanol gradients. Viral particles contained in the 40% iodixanol layer were collected dropwise into 2mL microcentrifuge tubes, combined and concentrated using Amicon Ultra-15 centrifugal filter units with a molecular weight cut-off of 100 kDa (Merck Millipore). Before the concentration step, Amicon membranes were covered with 15 mL of 0.1% Pluronic F68/PBS solution for 10 min, then discarded and replaced with 15 mL of 0.01% Pluronic F68 in PBS. Tubes were centrifuged at 3.000 rpm for 5 min at 4 °C, and the flow-through was discarded. 15 mL of 0.001% Pluronic F68 in PBS + NaCl 200 mM solution was added and centrifuged at 3.000 x rpm for 5 min at 4 °C and the flowthrough was again discarded. During the concentration process, a buffer exchange was performed with the formulation buffer (FB: 0.001% Pluronic F68 in PBS). FB was added to the AAV-iodixanol sample to replace iodixanol and to avoid toxicity in animals after AAV injection. Purified AAV virions were aliquoted and stored at −80°C. The viral titer was quantified using RT-qPCR.

### Transfection protocol

HEK293T cells were plated in 24-well plates the day before transfection on poly-L-lysine-coated (8.3 μM) 13 mm glass coverslips and maintained at 37°C in 5% CO_2_ and 95% O_2_. HEK293T cells were transfected with 250 ng of plasmid DNA expressing either mouse wild-type Trpa1 (TRPA1) or the mutated gene (TRPA1*), using Lipofectamine™ 3000 Transfection Reagent (#L3000001, Thermo Fisher) according to the manufacturer’s instructions.

### Electrophysiology

Whole□cell patch□clamp recordings were performed on 13 mm coated coverslips as previously reported^78,85^. Briefly, coverslips were transferred to a recording chamber (500 µl volume), mounted on the platform of an inverted microscope (Olympus CKX41, Milan, Italy) and superfused at a flow rate of 1.5 mL·min^−1^ with a standard extracellular solution containing (in mM): 10 HEPES, 10 D□glucose, 147 NaCl, 4 KCl, 1 MgCl_2_ and 2 CaCl_2_ (pH adjusted to 7.4 with NaOH). Cells were voltage-clamped at −60 mV by 2-4 MW-recording electrodes filled with the following intracellular solution (in mM): CsCl 120; Mg_2_-ATP3; EGTA10 and HEPES10 (pH 7.4 with CsOH). All experiments were performed at room temperature (RT: 20-22°C). Data were acquired with an Axopatch 200B amplifier (Axon Instruments, Union City, CA), low-pass filtered at 10 kHz, stored, and analysed with pClamp 9.2 software (Axon Instruments). Series resistance (Rs), membrane resistance (Rm), and membrane capacitance (Cm) were routinely measured using fast hyperpolarizing voltage steps of 10 mV. Only cells showing a stable Cm and Rs (<20% variation) throughout the experiment were included in the analysis. A Voltage ramp protocol (−100 to +100 mV; 800 ms) was applied every 15 s to evoke overall voltage-dependent currents in TRPA1- or TRPA1*-transfected HEK293T cells. After a stable baseline was established, AITC (100 µM) or 4-HNE (100 µM) was applied for 2 min, followed by the selective TRPA1 antagonist A967079 (100 µM). TRPA1- or TRPA1*-activated currents were obtained by subtraction of the ramp trace recorded after adding the TRPA1 antagonist A967079 from that recorded in the presence of each agonist. Where indicated, a voltage step protocol (−140 to +140 mV, 20 mV increment; 300 ms step duration) was applied before or during agonist exposure.

### H_2_O_2_ *in vitro* imaging

The genetically encoded probe for H_2_O_2_-HyPer [HyPer7.2^39^] was used on TRPA1 or TRPA1*-HEK293T transfected live cells. Briefly, cells were plated on poly-L-lysine-coated (8.3 μM) 35-mm glass coverslips and transfected with HyPer7.2 DNA (2 μg) using jetOPTIMUS® DNA transfection reagent (#55-250; Polyplus, Lexington, MA, USA). Cells were washed and transferred to a chamber on the stage of a fluorescent microscope for recording (Axio Observer 7; with a fast filter wheel and Digi-4 lens to record excitations; ZEISS, Stuttgart, Germany) after an incubation period of 24–48 h at 37C° with 5% CO_2_ and 95% O_2_. Depending on the experiment, cells were exposed hydrogen peroxide (H_2_O_2_, 3 nM-3 µM), AITC (10 nM-10 µM), 4-HNE (10 nM-10 µM) or vehicle (0.0001% DMSO) and H_2_O_2_ variations were monitored for approximately 300 s. EC_80_ for H_2_O_2_ (TRPA1= 809 nM, TRPA1*= 394 nM), AITC (TRPA1= 1.5 µM, TRPA1*=1.9 µM) and 4-HNE (TRPA1 = 4.4 µM, TRPA1* = 3.3 µM) was used in the presence of A967079 (30□μM) or vehicle (0.0001% DMSO). The ΔF/F0 _408/455_ ratio was calculated for each experiment, and the results were expressed as the area under the curve (AUC).

### Immunofluorescence

Anesthetized mice were transcardially perfused with PBS (phosphate-buffered saline), and then tissues (sciatic nerve and dorsal root ganglia, DRG) were collected. Tissues were fixed with 10% formalin overnight at 4°C, and then paraffin-embedded or cryoprotected in 30% sucrose overnight. Frozen samples were cryosectioned at 10µm while paraffin samples were cut at 5 µm with microtome. Frozen sections of mouse sciatic nerve were incubated with primary antibody 4-HNE (#ab48508, mouse monoclonal (HNEJ-2), Abcam, 1:25) diluted in fresh blocking solution normal goat serum (NGS, 5% in PBS-T) for 1 h at room temperature. Sections were then incubated for 2 h in the dark at room temperature with fluorescent secondary antibody polyclonal Alexa Fluor 647 (#A21236, goat anti-mouse polyclonal, Thermo Fisher Scientific, 1:600, RRID:AB_2535805) diluted in blocking solution, then washed and coverslipped using a water-based mounting medium with DAPI (#ab104139, Abcam). The 4-HNE staining was evaluated by measuring fluorescence intensity using an image processing program (ImageJ 1.32J, National Institutes of Health).

For formalin-fixed, paraffin-embedded sections, the primary antibody was used after antibody retrieval with citrate buffer (pH 6) and blocking for 1 h with fresh blocking solution containing normal donkey serum (NDS, 5% in PBS-T). Primary antibodies S100 (#ab196175, rabbit, 1:50), NeuN (#MAB377, mouse monoclonal, Merck, 1:400), and GFP-FITC conjugated (#ab6662, goat, 1:200) were diluted in NDS 5% and incubated overnight at 4°C. Alexa Fluor 647 (#A21236, goat anti-mouse polyclonal, Thermo Fisher Scientific, 1:600) diluted in NDS 5% was used as a secondary antibody, incubated for 2 h at RT. All slides were mounted with DAPI. All slides were visualized and analyzed using a Zeiss Axio Imager 2 microscope in Z-stack mode (Zeiss). The number of S100- or NeuN-positive cells expressing GFP was manually counted at 600X magnification. We acknowledge that detecting biomarkers localized in two or more intracellular compartments of the same cell may be affected by slice orientation, leading to incomplete colocalization, and that this is a limitation of the method.

### Southern blot

All Southern blots were performed using radioactively-labelled probes (α32P) as previously described^86^

### Electroporation and Selection of ESCs

Electroporation and screening of ESCs were performed as previously described^87^. Briefly, the targeting construct was electroporated using the Gene Pulser Xcell Electroporation System (#165-2661, Bio-Rad, Hercules, CA, USA). In brief, 10 μg of circular targeting vector was combined with 5 × 10^6 ESCs in a 4 mm electroporation cuvette and pulsed at 400 V, 125 μF. Cells were distributed on two 10 cm plates with DR4 feeders (mouse embryonic fibroblasts derived from the Dnmt1tm3Jae Hprtb-m3 Tg(pPWL512hyg)1Ems/J line; ^88^ in KSR medium. Twenty-four hours after electroporation, the medium was renewed with KSR containing G418 (150 μg/mL). This selection medium was changed every day until clone picking (10 days after electroporation). Picked colonies were trypsinized and triplicated into 96-well plates. One plate was used for the initial screen; the second plate was used to amplify the selected (Long Range-PCR-positive) clones, and the third plate was reserved as a backup. Cells were allowed to grow again in 96-well plates containing feeders for about 4–5 days in KSR medium.

#### Cell Lysis

Plates were washed with PBS. 75 μL of DirectPCR Lysis Reagent (#302-C Viagen Biotech) and 5 μL proteinase K (10 mg/mL) were added to each well. Plates were wrapped with foil lids and shaken slowly and incubated for 5 h to overnight on a plate shaker at 55°C. The next day, the plates were shaken for a few minutes to homogenize the lysate. 80 μL of H_2_O was added to each well to compensate evaporation. Three wells (one with normal, one with high and one with lower cell content) were kept apart and served as controls for the PCR. In a 96-well plate, 1 μL of each ESC lysate was added to 5 μL of a solution of Tris HCl at 5 mM, and proteinase K was inactivated 5 min at 95°C.

### Embryonic Stem Cell Screening and Validation

#### Long-Range PCR

The gene-specific LR-PCR genotyping primers used to identify clones were designed as follows: primers were selected by examining the genomic sequences flanking the 5′ and 3′ targeting vector homology arms, ensuring they avoided repeated sequences. The 3′ LR-PCR was performed with a gene-specific reverse primer (Rext; 5’-AACAATGTCACCTGCTTGCACTAGC-3’) and a forward selection cassette universal primer (Fneo; 5’-AGGGGCTCGCGCCAGCCGAACTGTT-3’).

The 5′ LR-PCR is also combined in a mixture with a universal reverse primer (Rneo; 5’-CTGCACGAGACTAGTGAGACGTGC-3’) with a gene-specific forward primer (Fext; 5’-GTGTAGGTGTTCAGATCTGTGTTCC-3’). The 5’ LoxP-PCR was performed with the same gene-specific forward primer (Fext; 5’-GTGTAGGTGTTCAGATCTGTGTTCC-3’) and a LoxP specific reverse primer (Rlox; 5’-CATACATTATACGAAGTTATGGCCG-3’).

Screening reactions were performed with a combination of two DNA polymerases: Red Hot Taq polymerase (previously ABGENE #AB-0406/B, special production by Thermo Fisher) and Pwo DNA polymerase (#11644955001, Sigma-Aldrich). Genotyping was carried out on 96 well plates using 20 μL reaction volumes. In each well except the 3 last ones was placed: 2 μL universal or gene-specific forward primer (1.25 μM), 2 μL gene-specific or universal reverse primer (1.25 μM), 6 μL DNA lysate (corresponding to approximately 50 ng), 2 μL 10× buffer, 1.2 μL MgCl_2_ (25 mM), 0.4 μL dNTP (10 mM), 0.15 μL Red Hot Taq polymerase (5 U/μL), 0.02 μL Pwo DNA polymerase (5 U/μL) and 6.23 μL water. To validate each assay, as no positive control clone was available, the last three wells were kept for control PCR designed as follows: the internal universal primer was replaced with a gene-specific primer (5’-GAGCTGCATGTGTGAATTAAATTCT-3’) that was designed at the extremity of the most distal part of the homology arm. The screen was considered valid if at least one of these wells showed a PCR fragment at the expected size. The cycling conditions were as follows: denaturation step of 5 min at 96°C, then 30 cycles of (96°C for 8 s, 60°C for 10 s, 68°C for 6 min) with an increase of 15 s at each cycle followed by 5 cycles of (96°C for 8 s, 60°C for 10 s, 68°C for 8 min); to finish, the samples were incubated at 68°C for 10 minutes.

### DNA Sequencing Library Preparation, and Analysis

Sciatic nerves were collected from mice and stored in RNA later at −80°C until processing. Total RNA was extracted with RNeasy® Lipid Tissue Mini kit (#74804; Qiagen SpA, Hilden, Germany) according to the manufacturer’s protocol. The RNA quantity and quality were determined by measuring the absorbance at 260/280 nm with QIAxpert System. RNA was reverse transcribed using SuperScript™ IV VILO™ Master Mix (Invitrogen) following the manufacturer’s protocol. *Wild Type*, CKI and mutated Trpa1 cDNA sequences from exon 18 to exon 24 were amplified with KOD Hot Start DNA Polymerase (#71086, Sigma Aldrich)(primers Fw 5’-TGTCATGGTCCAACATAACCGCA-3’; Rev: 5’-GCATGCTTCTGGACCTCAGC-3’). PCR products were purified using Gel and PCR Clean-up (NucleoSpin®; #740609.50, MACHEREY-NAGEL) according to the manufacturer’s instructions. PCR products were used to create a multiplexing library using Ligation sequencing amplicons - Native Barcoding Kit 24 V14 (SQK-NBD114.24, Oxford Nanopore Technologies) according to the manufacturer’s protocol. For each sample, 200 fmol of amplicon were used to prepare the library. The first step was end-prep using the NEBNext® Ultra™ II End Repair/dA-Tailing Module (NEB, E7546). This step was followed by barcode ligation and subsequent adapter ligation. The resulting libraries were multiplexed and sequenced on a PromethION 2 Solo device (Oxford Nanopore Technologies) using the MinKNOW software for 12 hours following manufacturer’s protocol.

### Bioinformatic analysis

Reads were filtered to retain those with a minimum quality score of 10 and a minimum length of 500 base pairs. After quality control (QC) filtering, reads were aligned using the minimap2 tool (version 2.17-r941) with the command option "-ax map-ont", to a custom reference containing the Mus musculus GRCm39 genome and the sequences of the cKI and CTL vectors. Aligned reads were converted to BAM and sorted with the SAMtools (version 1.10) tool. Coverage of the regions was calculated using mosdepth (version 0.3.12). Following alignment, reads were visualized using IGV software (version 2.19.6). Sequencing yielded a total of 104,360 reads for MiniG, 102,772 for GEdit, and 53,092 for cKI. The percentage of mapped reads was 72% for MiniG (75,797 reads), 71% for GEdit (73,257 reads), and 12% for cKI (6,201 reads). In the region of interest, approximately 6% of reads for both MiniG and GEdit showed the N858S mutation. Additionally, 24 reads were aligned to the cKI vector sequence.

### Statistical analysis

The results are expressed as the mean ± SEM. For multiple comparisons, a one-way ANOVA followed by a post hoc Bonferroni test was used. The two groups were compared using Student’s t-test. For behavioral experiments with repeated measures, a two-way mixed ANOVA followed by a post hoc Bonferroni test was used. Statistical analyses were performed on raw data using GraphPad Prism 8 (GraphPad Software Inc.). P-values less than 0.05 (P□<□0.05) were considered significant. EC_50_ values were determined from non-linear regression models using GraphPad Prism 8. The statistical tests used and sample size for each analysis are shown in the figure legends.

