## Supplementary material for "Disentangling Schwann Cell and Neuronal TRPA1 Function in Mouse Models of Familial Episodic Pain Syndrome": Supll Material

**Supplementary figure legends**

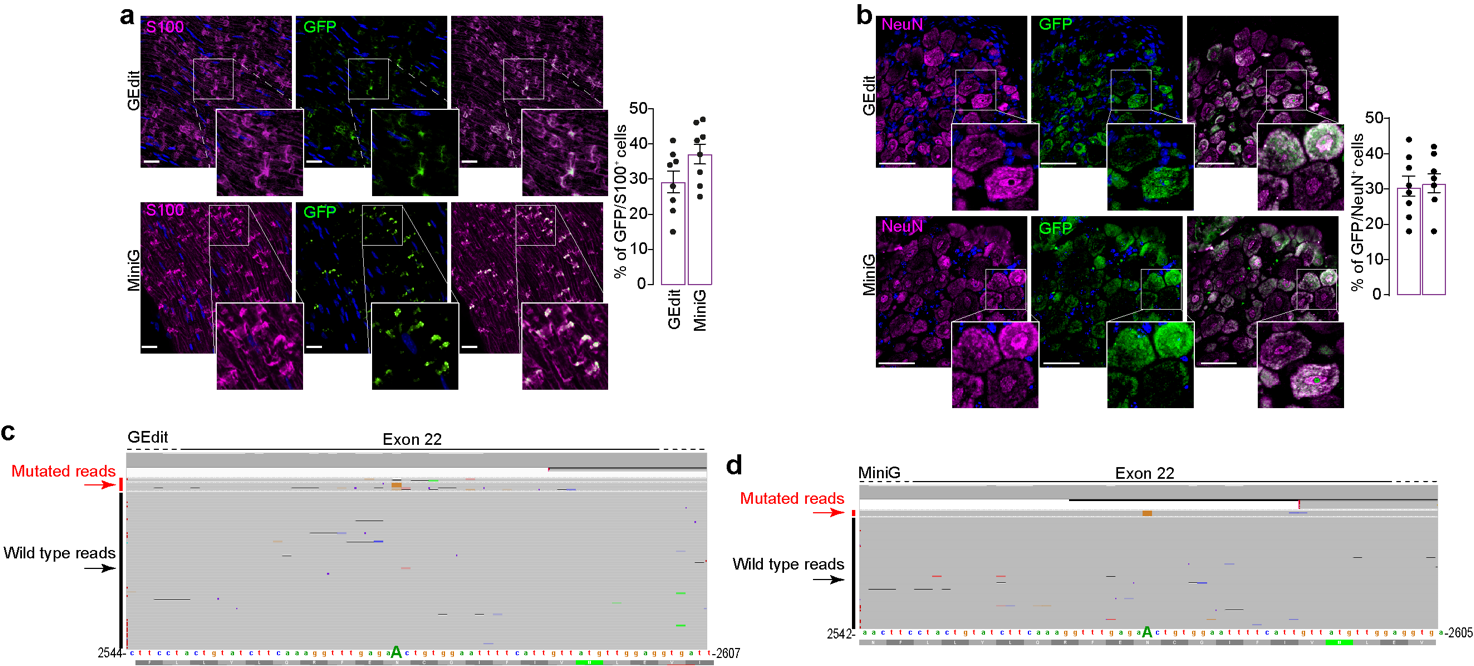

**Figure S1. *Trpa1** Nanopore sequence alignments in GEdit and MiniG mice.** Representative images and cumulative data (% of eGFP positive cells) in GEdit and MiniG approaches efficiency expressed as eGFP positive cells in **(a)** mouse sciatic nerve and **(b)** dorsal root ganglia primary sensory neurons (scale bar: 10 µm). eGFP expression is associated with saCas9 expression. Data are mean ± s.e.m. Integrated Genome Viewer (IGV version 2.19.6) visualization of long-read alignments from **(c)** GEdit and **(d)** MiniG mice. The reference sequence (color letters) represents the vector sequence, with the *Trpa1* wild-type base (A) shown in uppercase. Mutated base (G) for *Trpa1** is highlighted in orange.

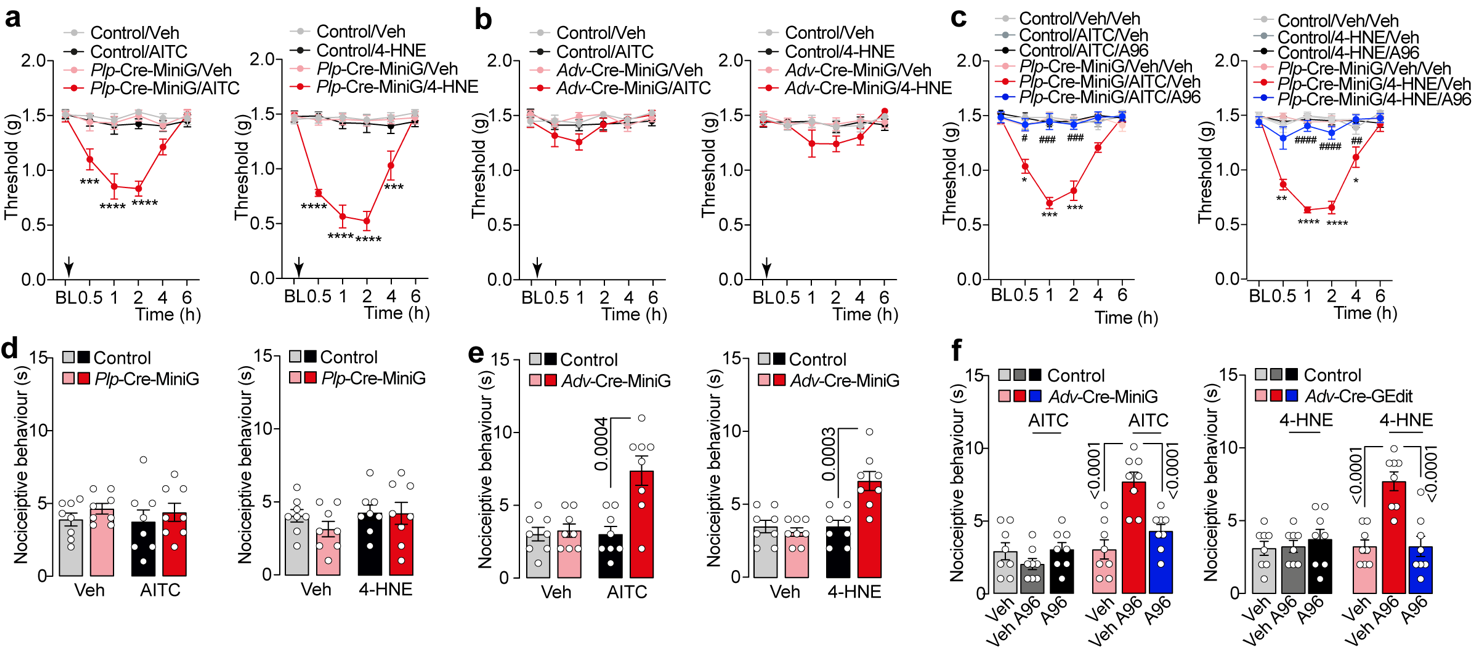

**Figure S2. Allodynia and non-evoked nociception in mouse with mutated TRPA1 in Schwann cells and primary sensory neurons.**

**(a,b)** Mechanical allodynia after intraplantar (i.pl./10 μl) injection of AITC (0.1 nmol), 4-HNE, (0.5 nmol) or vehicle (Veh) in *Plp*-Cre-MiniG, *Adv*-Cre-MiniG or Control mice. **(c)** Mechanical allodynia after i.pl. AITC (0.1 nmol), 4-HNE, (0.5 nmol) or Veh in *Plp*-Cre-MiniG mice pretreated with A967079 (A96, 100 mg/kg, intraperitoneal, i.p.) or Veh. **(d,e)** Acute nociception after i.pl. AITC AITC (0.1 nmol), 4-HNE, (0.5 nmol) or Veh in *Plp*-Cre-MiniG, *Adv*-Cre-MiniG or Control mice. **(f)** Acute nociception after i.pl. AITC (0.1 nmol), 4-HNE, (0.5 nmol) or Veh in *Adv*-Cre-MiniG, *Adv*-Cre-GEdit mice pretreated with A96 (100 mg/kg, i.p.) or Veh. Data are mean ± s.e.m. (n=8 mice/group). Arrow indicates time of compounds administration. 1-way and 2-way ANOVA, Bonferroni correction. *p<0.05, **p<0.01, ***p<0.001, ****p<0.0001 vs Control/Veh, Control/Veh/Veh; #p<0.05, ##p<0.01, ###p<0.001, ####p<0.0001 vs *Plp*-*Cre*-MiniG/Veh/Veh.

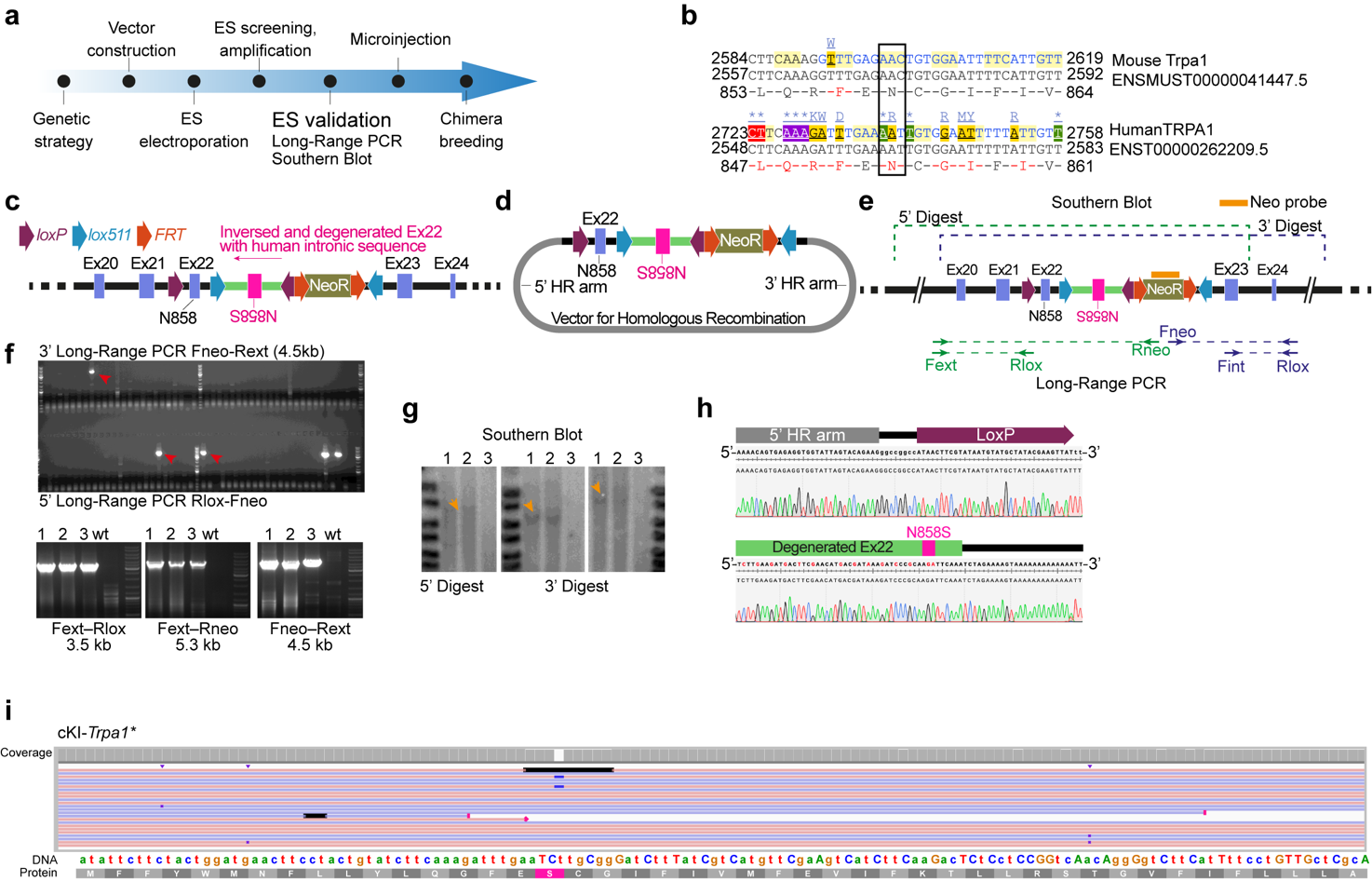

**Figure S3. Generation of a conditional knock-in (cKI) mouse model carrying a heterozygous FEPS mutation. (a)** Experimental timeline and workflow for development of the cKI mouse model. **(b)** Partial sequences of mouse *Trpa1* and human *TRPA1*, highlighting DNA triplet encoding Asparagine. **(c)** Schematic overview of the genetic strategy used to generate the FEPS cKI mouse model. **(d)** Targeting vector containing the cKI cassette flanked by homology arms for electroporation into mouse embryonic stem (ES) cells. **(e)** Schematic representation of 5′ and 3′ long-range PCR and Southern blot analysis using a Neo probe. **(f)** Screening of recombinant ES clones by 3′ long-range PCR (red arrow) and subsequent validation by 5′ long-range PCR of the three candidate clones identified. **(g)** Southern blot analysis following 5′ and 3′ enzymatic digestions using HindIII and SacI or PacI, respectively; orange arrows indicate the expected bands. **(h)** Validation of the 5′ *LoxP* site (top) and the inverted, degenerated exon 22 containing human intronic sequence by Sanger sequencing (bottom). **(i)** Integrated Genome Viewer (IGV version 2.19.6) visualization of long-read alignment from cKI mice. The reference sequence (color letters) represents the vector sequence, with degenerated bases shown in uppercase.

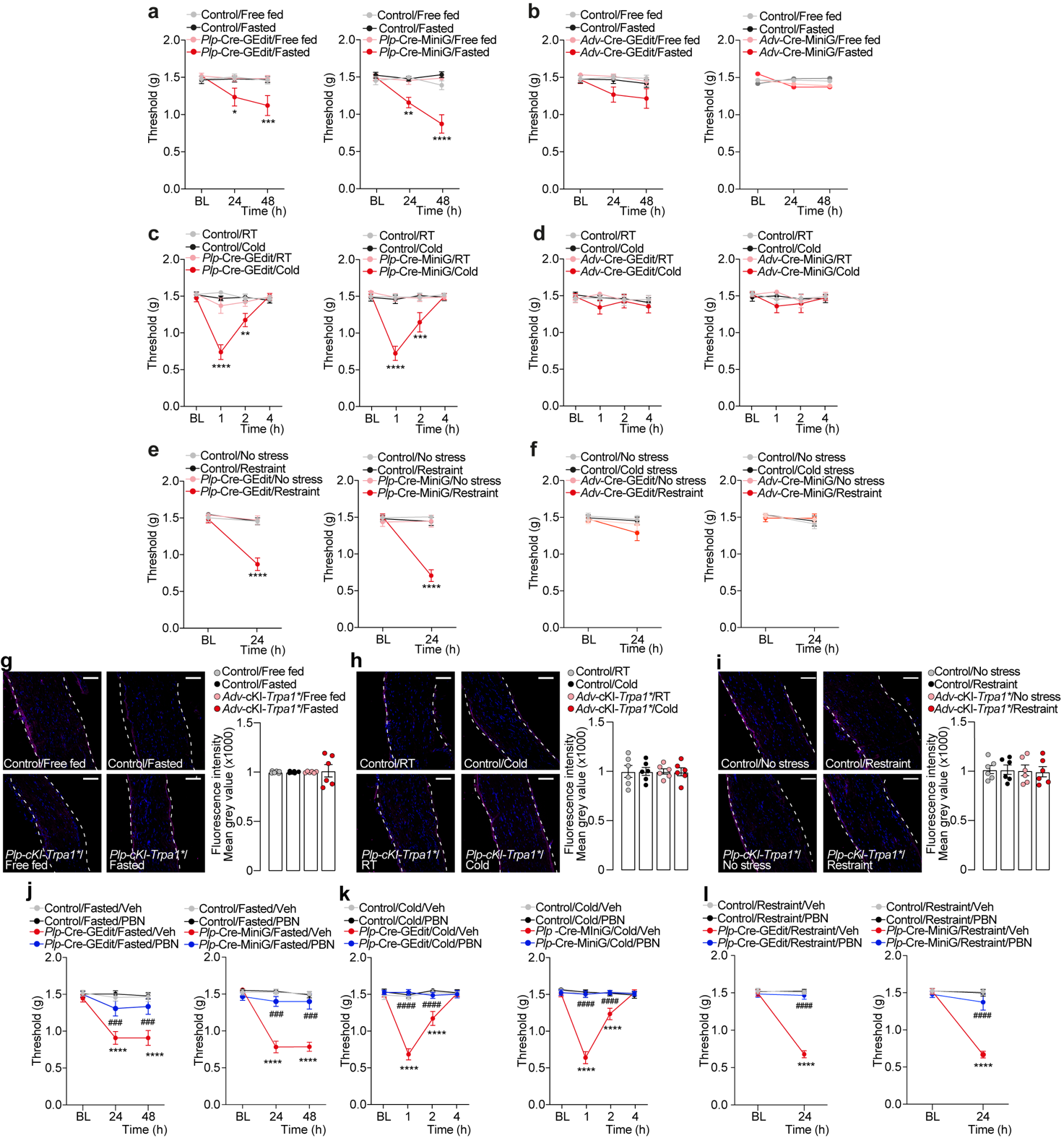

**Figure S4 (a,b)** Time dependent mechanical allodynia in Free fed or Fasted *Plp*-Cre-GEdit, *Plp*-Cre-MiniG, *Adv*-Cre-GEdit, *Adv*-Cre-MiniG or Control mice. **(c,d)** Time dependent mechanical allodynia exposed to cold or room temperature (RT) in *Plp*-Cre-GEdit, *Plp*-Cre-MiniG, *Adv*-Cre-GEdit, *Adv*-Cre-MiniG or Control mice. **(e,f)** Time dependent mechanical allodynia in the presence or absence of restraint stress in *Plp*-Cre-GEdit, *Plp*-Cre-MiniG, *Adv*-Cre-GEdit, *Adv*-Cre-MiniG or Control mice. Representative images and cumulative data of fluorescence in sciatic nerve tissues of *Adv*-cKI-*Trpa1** or Control mice **(g)** free fed or fasted **(h)** exposed to cold or RT or **(i)** in the presence or absence of restraint stress. (n=6, scale bar: 50 μm). Time dependent mechanical allodynia following repeated administration of phenyl-N-tert-butylnitrone (PBN, 100 mg/kg,intraperitoneal, i.p.) or vehicle (Veh) in *Plp*-Cre-GEdit, *Plp*-Cre-MiniG or Control mice **(j)** fasted, **(k)** exposed to cold or **(l)** restraint stress Data are mean ± s.e.m. (n=8 mice/group). 1-way, 2-way ANOVA, Bonferroni correction. *p<0.05, **p<0.01, ***p<0.001, ****p<0.0001 vs Control/Fasted, Control/Cold, Control/Restraint or vs Control/Fasted/Veh, Control/Cold / Veh, Control/Restraint/Veh; ### p<0.001, ####p<0.0001vs *Plp-*Cre-GEdit/Fasted/Veh, *Plp-*Cre-MiniG/Fasted/Veh, *Plp*-Cre-GEdit/Cold/Veh, *Plp-*Cre-MiniG/Cold/Veh, *Plp-*Cre-GEdit/Restraint/Veh or *Plp-*Cre-MiniG/ Restraint /Veh.
